# Histoimmunogenetics Markup Language 1.0: Reporting Next Generation Sequencing-based HLA and KIR Genotyping

**DOI:** 10.1101/014951

**Authors:** Robert P. Milius, Michael Heuer, Daniel Valiga, Kathryn J. Doroschak, Caleb J. Kennedy, Yung –Tsi Bolon, Joel Schneider, Jane Pollack, Hwa Ran Kim, Nezih Cereb, Jill A. Hollenbach, Steven J. Mack, Martin Maiers

**Affiliations:** National Marrow Donor Program, MN USA; HistoGenetics LLC, USA; University of California – San Francisco; Children’s Hospital & Research Center Oakland, Oakland, CA, USA

**Keywords:** HML, NGS, HLA, KIR, MIRING, data standards, genotyping

## Abstract

We present an electronic format for exchanging data for HLA and KIR genotyping with extensions for next-generation sequencing (NGS). This format addresses NGS data exchange by refining the Histoimmunogenetics Markup Language (HML) to conform to the proposed Minimum Information for Reporting Immunogenomic NGS Genotyping (MIRING) reporting guidelines (miring.immunogenomics.org). Our refinements of HML include two major additions. First, NGS is supported by new XML structures to capture additional NGS data and metadata required to produce a genotyping result, including analysis-dependent (dynamic) and method-dependent (static) components. A full genotype, consensus sequence, and the surrounding metadata are included directly, while the raw sequence reads and platform documentation are externally referenced. Second, genotype ambiguity is fully represented by integrating Genotype List Strings, which use a hierarchical set of delimiters to represent allele and genotype ambiguity in a complete and accurate fashion. HML also continues to enable the transmission of legacy methods (e.g. site-specific oligonucleotide, sequence-specific priming, and sequence based typing (SBT)), adding features such as allowing multiple group-specific sequencing primers, and fully leveraging techniques that combine multiple methods to obtain a single result, such as SBT integrated with NGS.

**Abbreviations:**

BRIDG: Biomedical Research Integrated Domain Group
CDISC: Clinical Data Interchange Standards Consortium
DaSH: Data Standard Hackathon
EMBL: European Molecular Biology Laboratory
ENA: European Nucleotide Archive
FDA: Food and Drug Administration
GL: Genotype List
HML: Histoimmunogenetics Markup Language
HLA: Human Leucocyte Antigen
IMGT: ImMunoGeneTics
ISO: International Organization for Standardization
LSDAM: Life Sciences Domain Analysis Model
KIR: Killer-cell Immunoglobulin-like Receptor
MHC: Major Histocompatibility Complex
MIRING: Minimum Information for Reporting Immunogenomic NGS Genotyping
NCI: National Cancer Institute
NGS: Next Generation Sequencing
NMDP: National Marrow Donor Program
OID: Object Identifier
SBT: Sequence Based Typing
SSO: Sequence Specific Oligonucleotide
SSP: Sequence Specific Primer
URI: Uniform Resource Identifier
XML: eXtensible Markup Language

## 1. Introduction

Human leukocyte antigen (HLA) genotyping is fundamental for research and clinical practice in immunogenetics and histocompatibility. Methods for generating this information have dramatically improved in the last three decades, from pioneering serological methods to modern DNA-based genotyping methods [1], revealing over time a seemingly never ending expansion of allele diversity [2], and presenting the subsequent challenge of reinterpretation of earlier results in light of new knowledge [3–6]. It has become increasingly clear that in order to ‘future-proof’ the results as much as possible, data standards are needed for recording and reporting of these results which include not only the allele assignments, but also the metadata (data describing other data) surrounding laboratory methods and allele assignment rules [5,7].

A new generation of sequencing methods has compounded this challenge. These methods are characterized by massively parallel technologies leading to high throughput sequencing, allowing for routine interrogation of not only select regions of a gene, but also whole-exon or whole-gene sequencing. While these technologies, commonly known as next-generation sequencing (NGS), have existed for several years, their application to HLA has been challenging due to a high degree of allelic polymorphism, the lack of robust genomic reference alignments for the MHC region, and the complex metadata required for allele assignment and later reinterpretation as reference sequences are refined and expanded. These considerations emphasize the need for recording and reporting complete metadata surrounding data collection, and in particular for data processing and interpretation.

Recently, a group of histocompatibility and immunogenetics stakeholders including clinicians, researchers, instrument manufacturers and software developers has gathered in a series of meetings to develop standards for recording and reporting NGS based genotyping of HLA [8]. One of the goals of these meetings has been to identify the Minimum Information for Reporting Immunogenomic NGS Genotyping (MIRING) [9] based on the principles of the reporting guidelines for the Minimum Information for Biological and Biomedical Investigations (MIBBI) [10]. The MIRING identifies eight principles for reporting NGS based genotyping of immunogenomics data. While the MIRING provides principles and guidelines, it does not provide a technical specification.

To develop a technical specification that meets these principles, we have extended the Histoimmunogenetic Markup Language (HML) [11]. HML is an electronic messaging format based on XML and developed as a community standard allowing explicit representation of genotyping results for polymorphic immunogenetic gene systems such as HLA and KIR. Primary data in the form of Sequence Specific Oligonucleotide (SSO)/Sequence Specific Primer (SSP) reactivities and Sequence Based Typing (SBT) sequences can be linked with nomenclature-based allele assignment. In extending HML, we added the ability to include NGS based genotyping, including metadata surrounding the primary data and acquisition methods, consensus sequences, variant calling including novel polymorphisms, and reporting full allele and genotype ambiguity through the use of Genotype List (GL) Strings [12].

## 2. Methods

The development of HML 1.0 was led by the National Marrow Donor Program (NMDP) through a series of meetings and discussions with the HLA Information Exchange Data Format Standards Group, and later with the Immunogenomic NGS Data Consortium [8], a community of registries, clinical and research laboratories, and industry partners focused on identifying and addressing specific data-reporting requirements for NGS-based genotyping, and occurred in parallel with development of MIRING.

In September 2014, a community of stakeholders including researchers, clinicians, software developers, sequencing platform vendors, HLA and KIR sequence database developers and administrators, registry and donor center IT professionals, and laboratory support staff gathered to attend a Data Standards Hackathon (DaSH) [13] to develop new ways to exchange NGS data for HLA and KIR. During this event the current state of HML was vetted by the community, and at the same time MIRING was further refined. These exercises led to new substantial requirements for the final specification for HML 1.0.

A rapid prototyping approach was used for the development of XML schemas for HML. As schemas were developed, example messages were created with various scenarios of HLA and KIR genotyping and syntactically validated using automated validation tools. The schema and examples can be found at https://bioinformatics.bethematchclinical.org/HLA-Resources/HML/.

In designing HML, it was important to accommodate the use of pointers to external references and datasets. This allows users to reuse previously registered metadata, and to reference datasets that are impractical to include in the HML message. These include registered methodologies, reference sequences, and raw sequence reads.

## 3. Results

HML 1.0 retains similar overall structure to previous versions, but with notable changes. As with earlier versions, reporting of primary data is separated from allele assignment. NGS is supported by new XML structures to capture all NGS data and metadata required to produce a genotyping result, including analysis-dependent (dynamic) and method-dependent (static) components. Pointers to external locations refer to registered methods, raw NGS reads, and reference standards. A separate component describing consensus sequences and variants was created specifically to accommodate NGS data, but could be used for other methodologies if desired. This component includes metadata describing the consensus sequences such as references sequences, phasing information, expected copy number, sequence block continuity, and other metadata. Reporting of allele assignment with full genotypic and allelic ambiguity is achieved through the use of GL Strings.

The overall HML 1.0 structure is seen in Figure 1. An HML message has four main sections: the document header, the typing methods, the allele assignment, and the consensus sequence. An HML message will contain the document metadata, and data describing one or more samples, generated with one or more typing methods for one or more gene families.

**Figure 1.**
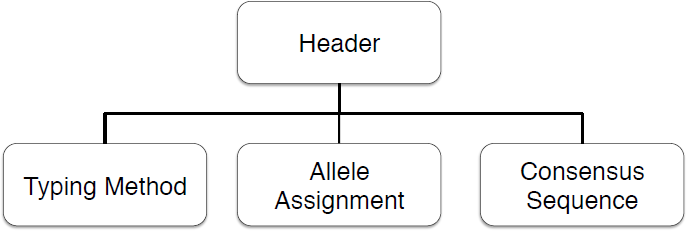
HML overall structure

In addition to several minor improvements and many other components left to retain backward compatibility, two major improvements were introduced in HML 1.0. First, new elements that conform to the MIRING reporting guidelines were added. Second, GL strings were added to allow expression of full allelic and genotypic ambiguity in addition to, and eventually in lieu of, NMDP allele codes. Although the expectation is that NGS will lead to unambiguous allele assignment through full gene sequencing, earlier methodologies will continue to be used and reported in HML 1.0, and in the near term NGS based genotyping may be limited by some laboratories to interrogating specific exons leading to allele and genotype ambiguity. Additional changes include enhancements to Sanger-sequence based typing (SBT-Sanger), external references to typing kit information, and expansions to include multiple gene families (e.g., HLA, KIR).

In the specification below, note that the declared element and attribute names are in Courier font to highlight their abstract nature.

### 3.1. HML Header

The header details are seen in Figure 2. Here, the message is contextualized, providing details such as identifying the lab sending the message, the report identifier, the sample(s) the report describes, and other reference information. This header does not contain details about any particular test, only the context in which it occurred.

**Figure 2.**
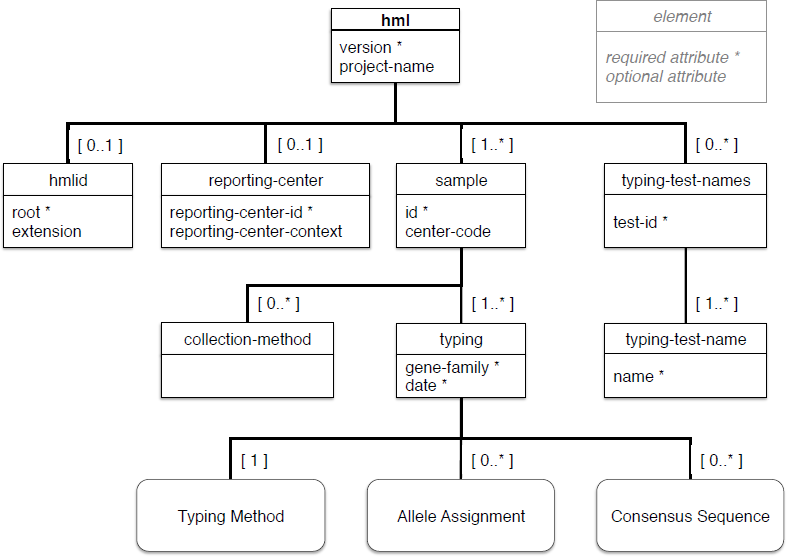
Header

#### 3.1.1. hmlid

A unique identifier for the HML document can be included through the hmlid element. The hmlid follows the HL7 convention for instance identifiers using a two-part key [14]. An ISO Object Identifier (OID) [15] may be provided through the `root` attribute and often represents the unique organization identifier publicly registered for an organization. The `extension` attribute contains the unique document id managed internally by the reporting organization. Together, the `root` and `extension` attributes combine to provide a globally unique identifier.

Alternatively the ISO OID may be extended directly with the `extension` in the `root`, or a Universally Unique Identifier (UUID), also known as Globally Unique Identifier (GUID), may be provided in the root without an `extension`. Examples of these different ways of using hmlid are found in Figure 3.

**Figure 3.**
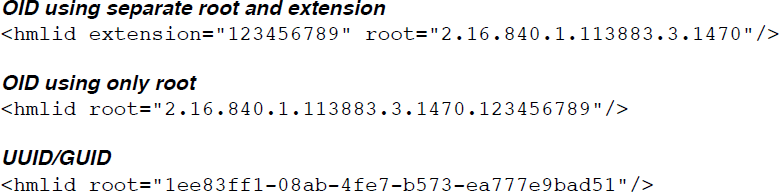
Examples of how hmlid can be used

#### 3.1.2. reporting-center

The `reporting-center` element identifies the entity/organization sending the HML message. If included, it must contain a unique `reporting-center-id` identifying the sender. A new attribute, `reporting-center-context`, has been added. This is used to report the context/naming authority of the identifier. This can be used by different organizations to identify the reporting organization using registered contact information. For example, although this element is optional, NMDP requires the `reporting-center` element in an operational context. If the reporting-center-context is not included, it is assumed to be “NMDP” and is a unique identifier that is registered with the NMDP. Another example of the use of this element is to refer to a testing lab registered in the NCBI Genetic Testing Registry (GTR) [16] through the GTR Lab ID. Figure 4 shows examples of how reporting-center-id may be used.

**Figure 4.**
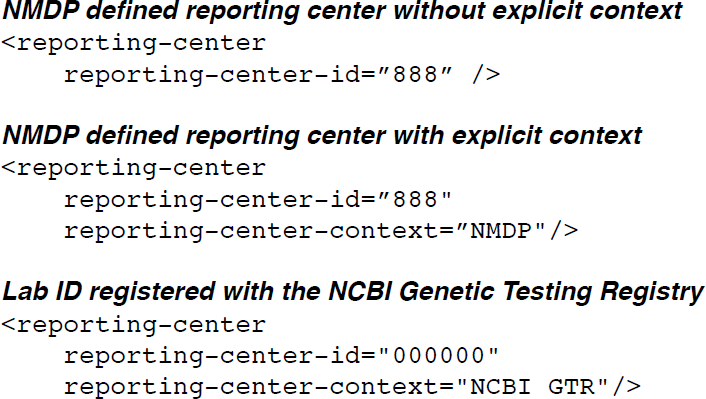
Examples of how reporting-center can be used

#### 3.1.3. typing-test-names

The `typing-test-names` element specifies a list of test names internally referenced by the SSO or SSP typing methods. It contains a `test-id` attribute associated with one or more `typing-test-name` containing test identifiers, which together are used to define a typing kit. It wraps a list of `typing-test-name` elements containing the test identifiers. The `test-id` is then used as an internal reference identifier to be used in the SSO and SSP typing methods sections.

#### 3.1.4. sample

An HML message contains one or more sample elements enclosing the genotyping data pertaining to a particular sample. The `id` and `center-code` attributes identify the sample and the organization where the sample originated (e.g., transplant or donor center) respectively. Multiple samples from the same individual, or different individuals can be included in a single HML message. A `collection-method` element may be used to identify how the sample was collected, e.g., swab, filter paper, blood aliquot. Each `sample` may be associated with multiple `typing` elements.

##### 3.1.4.1. typing

The `typing` element encapsulates the primary data from a genotyping method with a genotyping result (`allele-assignment`) that was determined from the primary data (`typing-method`) and/or consensus sequences (`consensus-sequence`). The `gene-family` attribute is used to constrain the typing to gene family name recognized by the HUGO Gene Nomenclature Committee [17] (e.g., HLA, KIR).

### 3.2. Genotyping Method and Primary Data

Figure 5 describes typing method and primary data elements. Here the method used to perform the genotyping and the primary un-interpreted results from that method are reported. This can be one of SSO, SSP, SBT-Sanger, or SBT-NGS.

**Figure 5.**
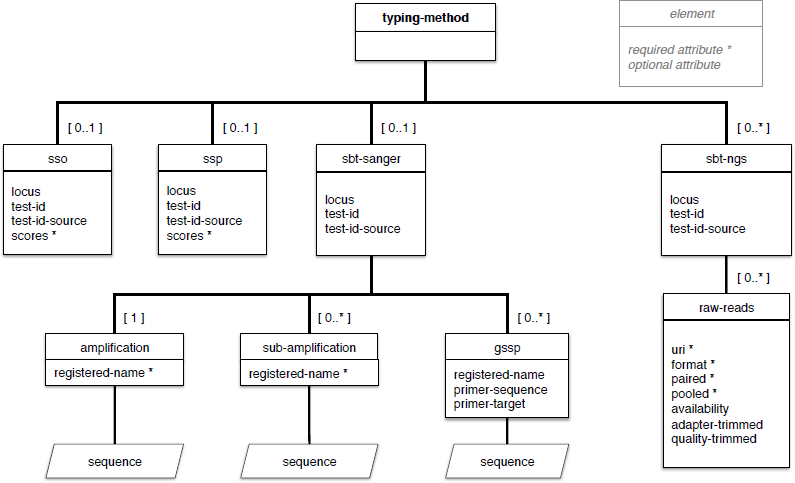
Typing methods & primary data

#### 3.2.1. Test IDs & Locus

To comply with MIRING requirements, three new attributes referencing additional metadata are available to more fully describe the method and locus targeted for each typing method. Two of these attributes name a test registrar (`test-id-source`) and an identifier (`test-id`) that can be dereferenced through the registrar. In previous versions of HML, sso and ssp elements referenced test information in the `test-id` element found in the header, which is used for encapsulating the test methods consisting of one or more individual tests or probe kits within the HML document. These new attributes allow labs to register their test method in an external testing registrars, such as the NCBI GTR [16].

The third new attribute for each typing method element is `locus`, which indicates the target of the typing method. `locus` is an optional attribute. If present this must be a gene name recognized by the HUGO Gene Nomenclature Committee [17], e.g., HLA-A, HLA-DRB1, KIR2DL1.

#### 3.2.2. sso & ssp

The `sso` and `ssp` elements describe SSO and SSP genotyping methods, respectively. These elements are unchanged from previous versions of HML except for the addition of `test-id`, `test-id-source`, and `locus` attributes (see above). For submissions to NMDP, a corresponding `typing-test-names`/`typing-test-name` structure found in the header is expected.

#### 3.2.3. sbt-sanger

The `sbt` (Sequence-Based Typing) element found in previous versions of HML has been renamed to `sbt-sanger`. This reflects the recognition that both Sanger and NGS typing methods are sequence based, but with differences in the metadata needed to support them. Other changes include an additional child element to support sub-amplification primers and results. These primers are used to resolve ambiguities and may be used either concurrently with or after the amplification step. In addition, multiple `GSSP` elements (Group Specific Sequencing Primers), describing PCR primer sequences used to amplify polymorphic regions of sequences, are now allowed. In previous versions of HML, only a single `GSSP` was allowed.

#### 3.2.4. sbt-ngs

The new `sbt-ngs` element describes Sequence Based Typing using high throughput methods, also known as NGS (next-generation sequencing). As with other methods, `test-id`, `test-id-source`, and `locus` attributes describe the method and the target locus of the test. Each `sbt-ngs` may contain multiple `raw-reads` child elements.

The `raw-reads` element references the raw sequence reads generated by an NGS platform. This data can be quite large even for relatively small regions of the genome. Because of this, if the `raw-reads` are provided, this information is not directly embedded in the HML document, but instead is referenced through a `uri` (Uniform Resource Identifier) attribute that points to an external location and a `format` attribute that describes how the data is stored (e.g., the NCBI Sequence Read Archive (SRA)). Other attributes include the availability of the raw reads (public, private, or with permission) and processing metadata indicating whether or not the data are pooled (`pooled`), if the adapters have been trimmed (`adapterTrimmed`), or whether low-quality sequences have been trimmed (`qualityTrimmed`).

### 3.3. Allele Assignment

Figure 6 describes the Allele Assignment section. Several changes from previous versions of HML were made here. The element that was previously named `interpretation` is now renamed `allele-assignment` to more accurately reflect its purpose. Also, this element includes new attributes that describe the allele database or other source of the nomenclature (`allele-db`, e.g., IMGT-HLA Database) and release version of the database (`allele-version`, e.g., 3.19.0) that was used for the allele assignment.

**Figure 6.**
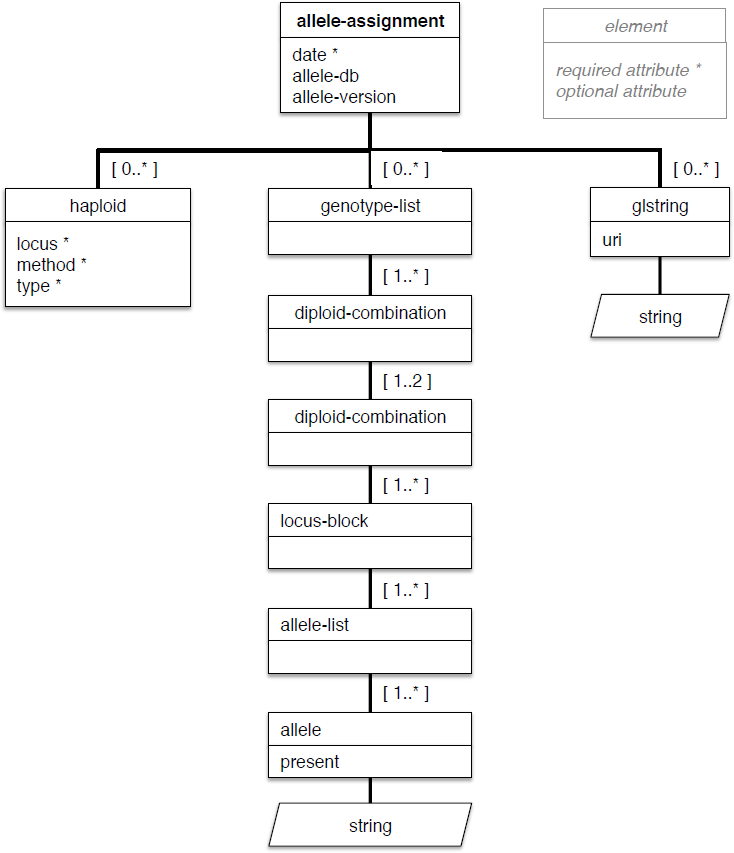
Allele Assignment

Allele assignment may be reported in three different formats. As in previous versions of HML, `haploid` and `genotype-list` elements are available and unchanged. A new third element representing Genotype List (GL) Strings (`glstring`), which allows allele assignments with full genotype and allele ambiguity [12], has been added. The GL String may be directly included, or a URI resource identifier pointing to a service that provides the GL String (e.g., https://gl.nmdp.org) may be used. An example of how a GL String may be reported is seen in Figure 7.

**Figure 7.**
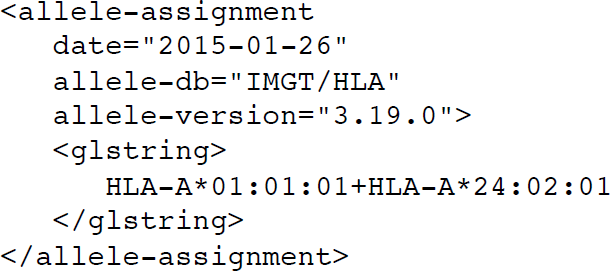
Example of how allele-assignment may be used with glstring

## 3.4. Consensus Sequence

Figure 8 describes the Consensus Sequence section. This is a new structure in HML that was created to provide the result of an alignment or assembly of shorter sequence reads generated by a NGS platform. It is dependent on a number of dynamic factors such as the processing pipeline including software and inputs, and references sequences used for alignments. This is not considered to be primary data, but rather as a type of interpretation. For this reason, it is not included within the typing method section, but is considered separately. While this section was created with NGS in mind, it may be used by other sequence-based typing methods if desired.

**Figure 8.**
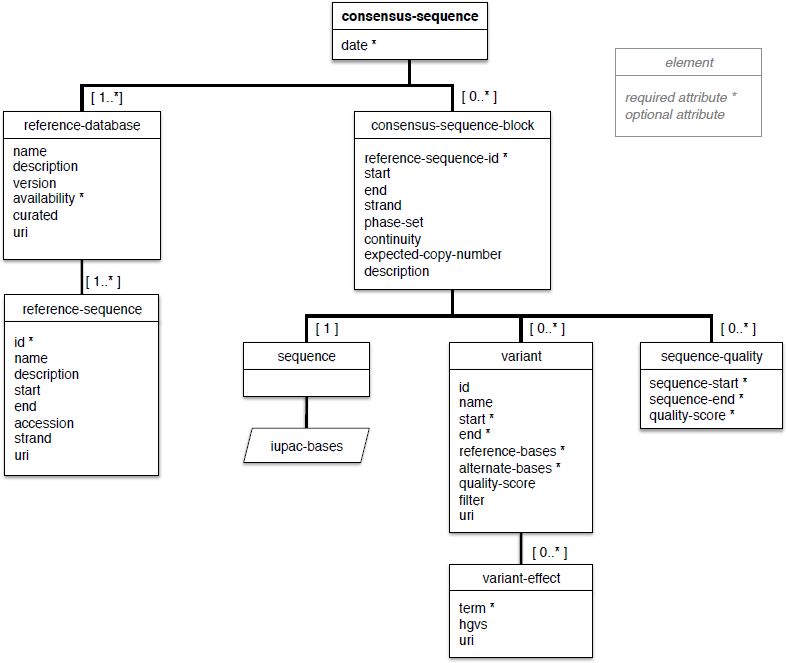
Consensus sequence

This section is largely based on many of the data elements found in the Variant Call Format (VCF) [18] and metadata identified from the MIRING [9]. Note HML uses `start` and `end` attributes with a 0-based coordinate system with closed-open ranges, *i.e*., the range includes the `start` position, but excludes the `end` position. This convention is used in many genomic data models (e.g., Global Alliance for Genomics and Health schemas [19,20], UCSC browser [21], BED files) because of its programming benefits. In contrast, the VCF specification indexes the 1st base having position 1.

A sequence is reported directly through the `sequence` element. A pointer to an external reference may also be included, or as variant of a reference sequence. If a variant is reported, the structure provides a method to report variant effects. Position specific quality values may also be recorded.

### 3.4.1. reference-database & reference-sequence

Together, the `reference-database` and `reference-sequence` elements describe the database containing the reference sequence used within the `consensus-sequence-blocks`. The `reference-database` may point to whole genome builds (e.g., Genome Reference Consortium), allele databases (e.g., IMGT-HLA Database), or another database containing sequences that are used as a reference (e.g., GenBank, EMBL-ENA). Metadata surrounding the `reference-database` include the name, a description, the version, its availability, whether or not the database is curated, and a URI pointing to its external location.

For each `reference-database`, one or more `reference-sequence` elements may be described. An id is used to uniquely identify the `reference-sequence` and is referenced by the `consensus-sequence-block`. Other metadata that may be provided as attributes include the name of the sequence, the start and end positions, an accession id, and a URI pointing the sequence directly.

Examples of how `reference-database` and `reference-sequence` can be used are found in Figure 9.

**Figure 9.**
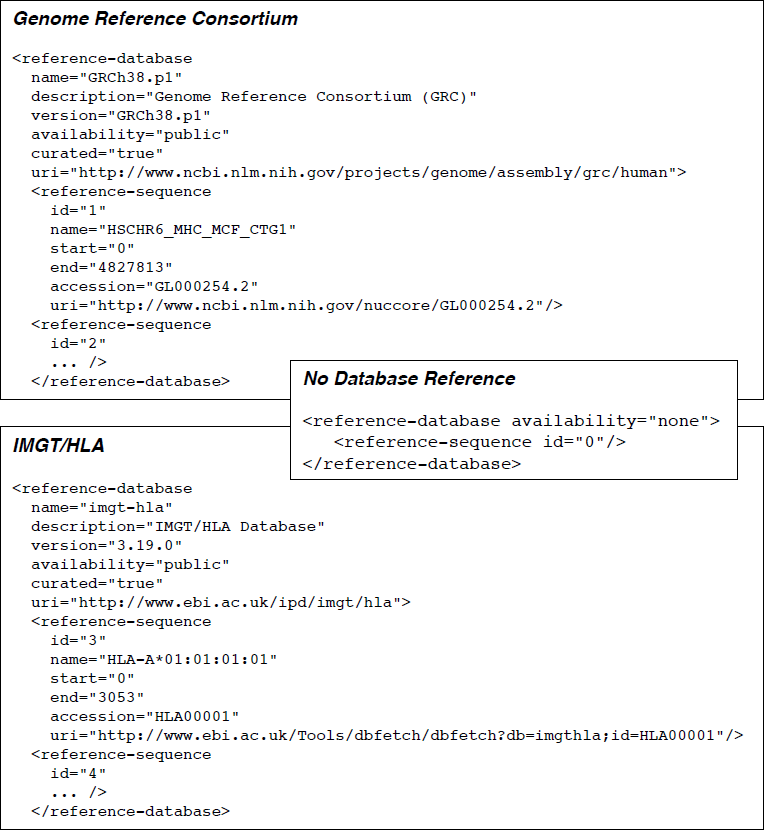
Examples of using reference-database and reference-sequence

### 3.4.2. consensus-sequence-block

A `consensus-sequence-block` encapsulates a DNA sequence consisting of IUPAC nucleotides [22] and variants against a reference sequence if an exact match is not found, and associated metadata. A `reference-sequence-id` is required and points to a `reference-sequence` found within the HML document. Start and end positions identify a targeted region on the reference. Other optional metadata are provided through attributes that describe the strand being identified, a description of the target region (e.g., “HLA-A exon 3”), a phase set identifier to associate different `consensus-sequence-blocks`, a continuity flag to indicate whether a gap exists between blocks, and an expected-copy-number to indicated how many copies of the sequence block was expected.

A quality score for positions within the consensus sequence may be captured in the `sequence-quality` element, using `start` and `end` attributes indicating the region of interest. Quality metrics for sequence quality vary by platform reflecting differences in sequencing chemistry and digital processing. Additional metrics, such as coverage depth at the sequence- and variant-levels (see below), lack common standards and may be calculated differently depending on the analytical method chosen. For these reasons HML quality reporting does not impose strict specification criteria.

### 3.4.3. variant & variant-effect

A `variant` element must be included if the `sequence` does not completely match the `reference-sequence`. A typical use for this element is describing a novel allele. The available attributes for `variant` corresponds with many data elements used in VCF files [18] and in the MIRING. The `id attribute` is used to uniquely identify the variant within the HML document. The `start` and `end` identify the coordinates of the reference sequence where the variant occurs. The nucleotides found at this region of the reference and what they are replace with are identified with the `reference-bases` and `alternate-bases` attributes, respectively.

An alternative external reference for the variant may be reported using the `uri` attribute. This could be a location for a VCF file, or an alternative format that represents the same information.

A child of `variant`, the `variant-effect` element reports the effect of this variation. The effect or effects should be described using Sequence Ontology (SO) variant effect terms [23] (e.g., missense_variant, stop_gained, downstream_gene_variant). Additional attributes may be used to provide more information on the effect, e.g., severity, POLYPHEN prediction, SIFT score [24,25]. In addition to the `term`, a Human Genome Variation Society (HGVS) nomenclature string [26,27] may be included. A `uri` may be used to provide an external reference for this variant effect. An example of how variants may be captured is seen in Figure 10.

**Figure 10.**
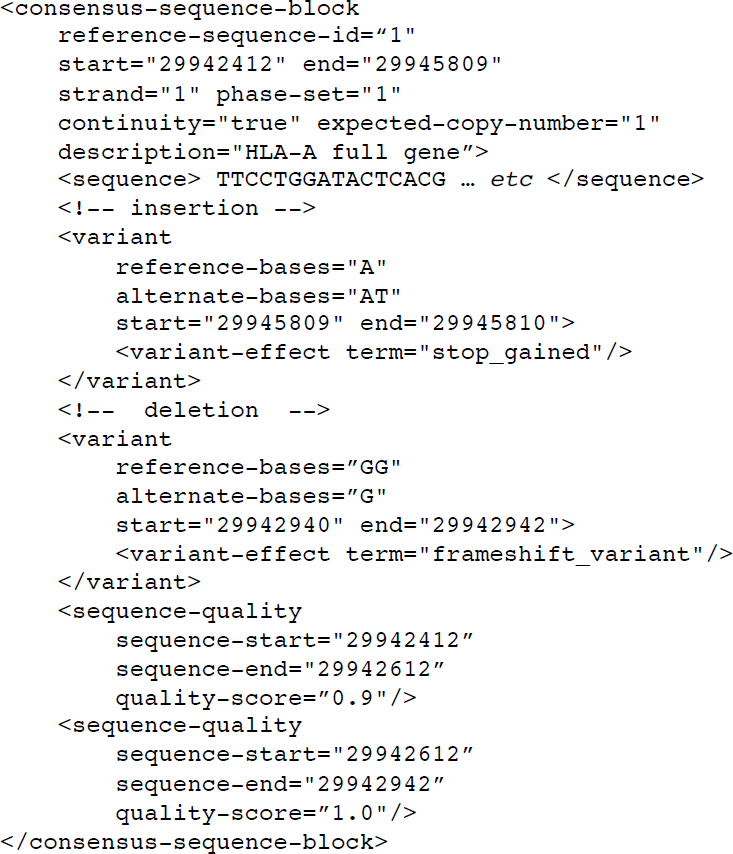
Example of consensus-sequence-block with variants

## 3.5. Extensibility

HML provides for extensibility using two methods. One is through the use of the `property` element that has been introduced as an optional child element for several nodes (`hml, sample, typing, typing-method types`, and `allele-assignment`). Here, name/value pairs (jointly defined and agreed upon by the creator and consumer of the HML document) may be entered. Additionally, `property` elements may contain custom nested structures of the creator’s choosing to add additional information and details to the enclosing element. The other method of extending the HML specification is through the use of the `anyAttribute` type associated with the `“consensus-sequence-block”`, `“sequence”`, `“variant”`, and `“variant-effect”` elements that allows additional attributes not specified by the schema.

## 4. Discussion

We present HML 1.0, an electronic format for exchanging data for genotyping of histoimmunogenetic markers such as HLA and KIR, with extensions for next-generation sequencing. These improvements equip the message with the mechanisms to collect new forms of typing data and accurately report genotype and allele ambiguity, in a machine-readable format that tightly pairs the typing method and result for downstream analyses.

## 4.1. MIRING

We have extended and enhanced HML to implement the principles and guidelines outlined by the MIRING. Eight elements were identified that need to be addressed in an MIRING compliant genotyping report.

MIRING elements and corresponding example solutions in HML 1.0 are presented in Table 1. In several cases, the MIRING offers technical solutions for how specific elements may be addressed, but these translate into different solutions in HML. For example, the pipe-delimited descriptor found in MIRING is required for FASTA files. Often this same information is found within the XML structure of HML and may not be explicitly needed. For example, in HML the `consensus-sequence-blocks` are captured sequentially, and each block contains within it all pertinent information within the attributes and child elements. Because of this, a separate identifier for the consensus sequence block to associate sequences with consensus sequences, references sequences, variants and associated metadata is not needed.

**Table 1.**
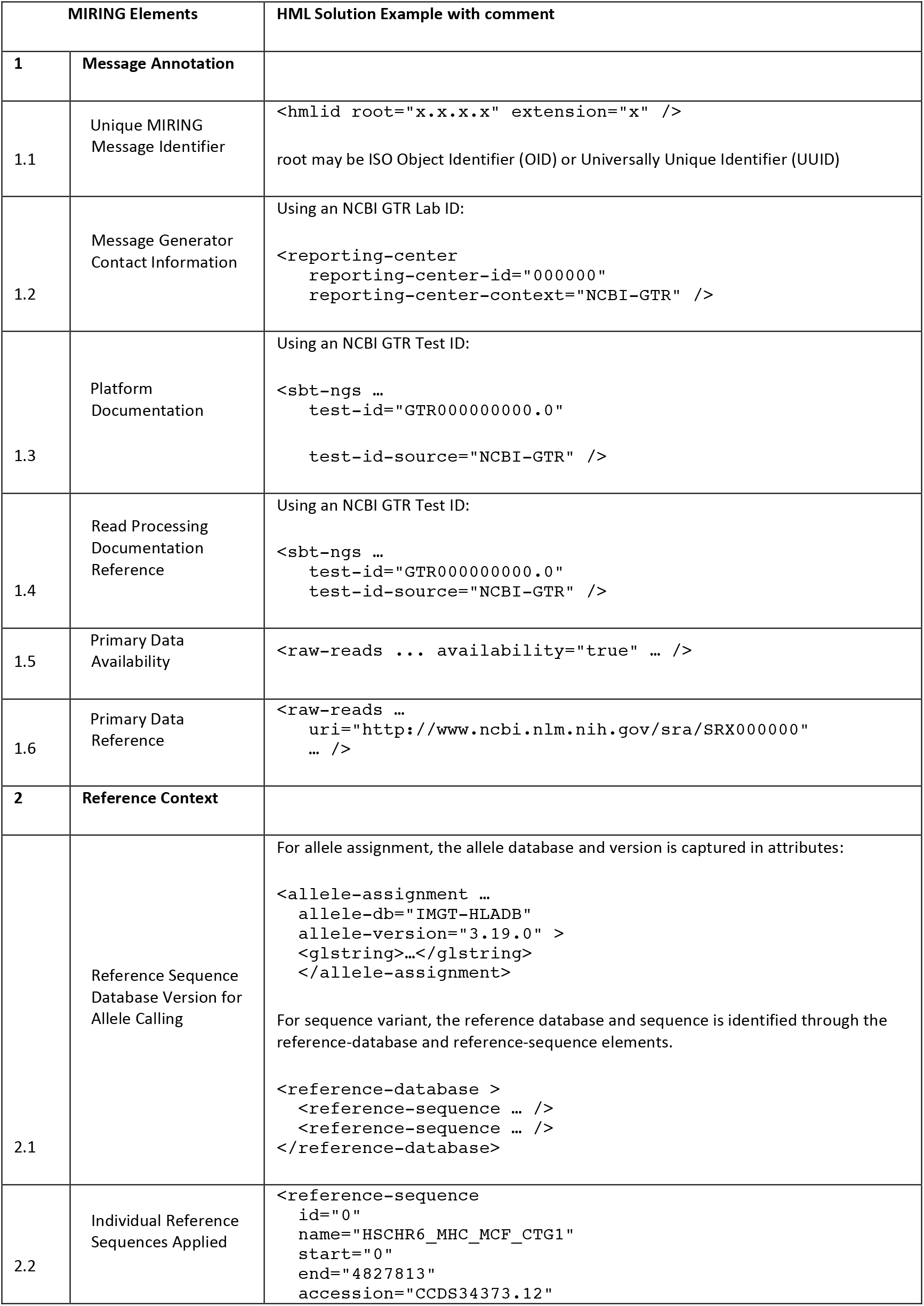
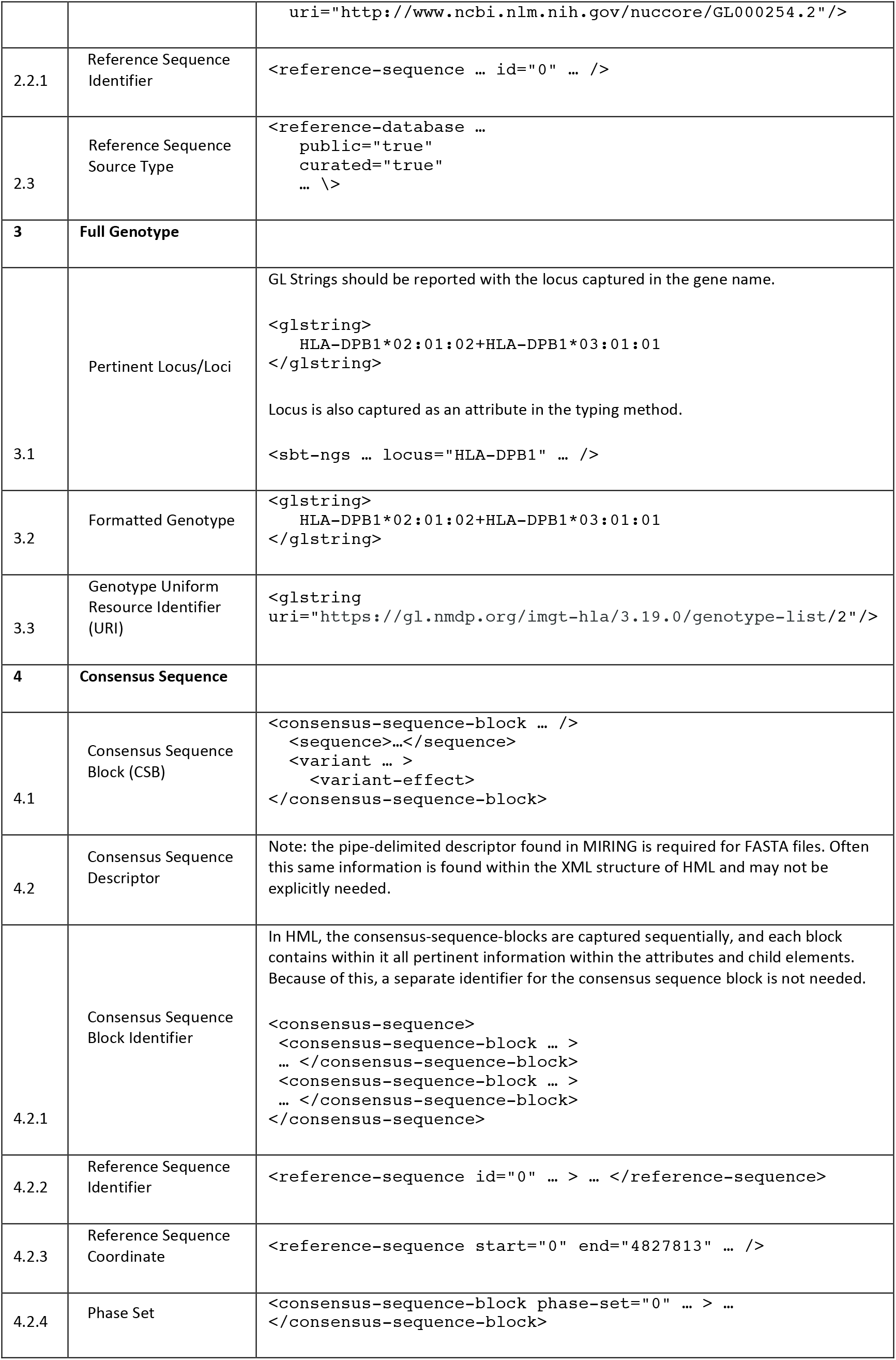
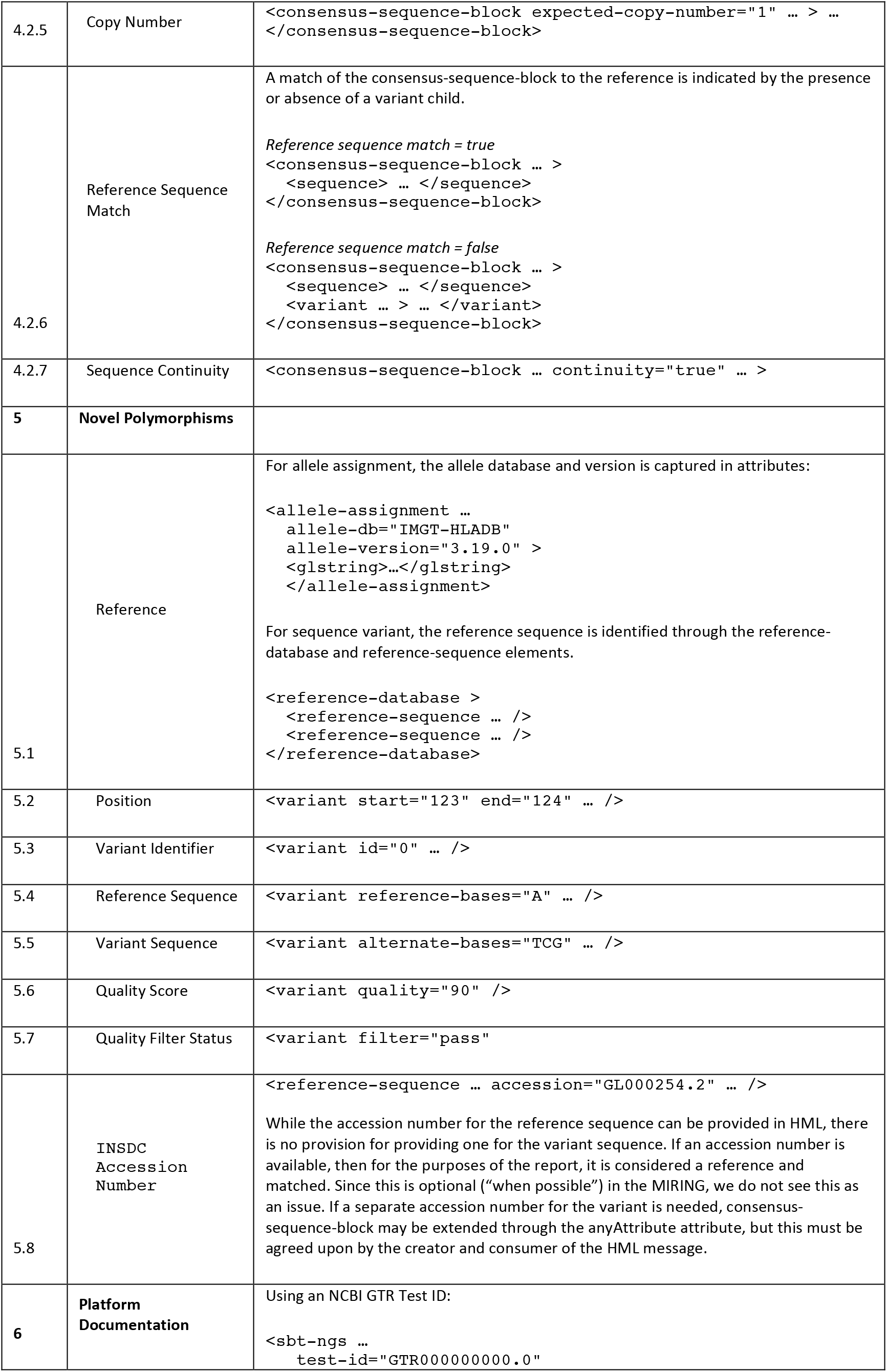
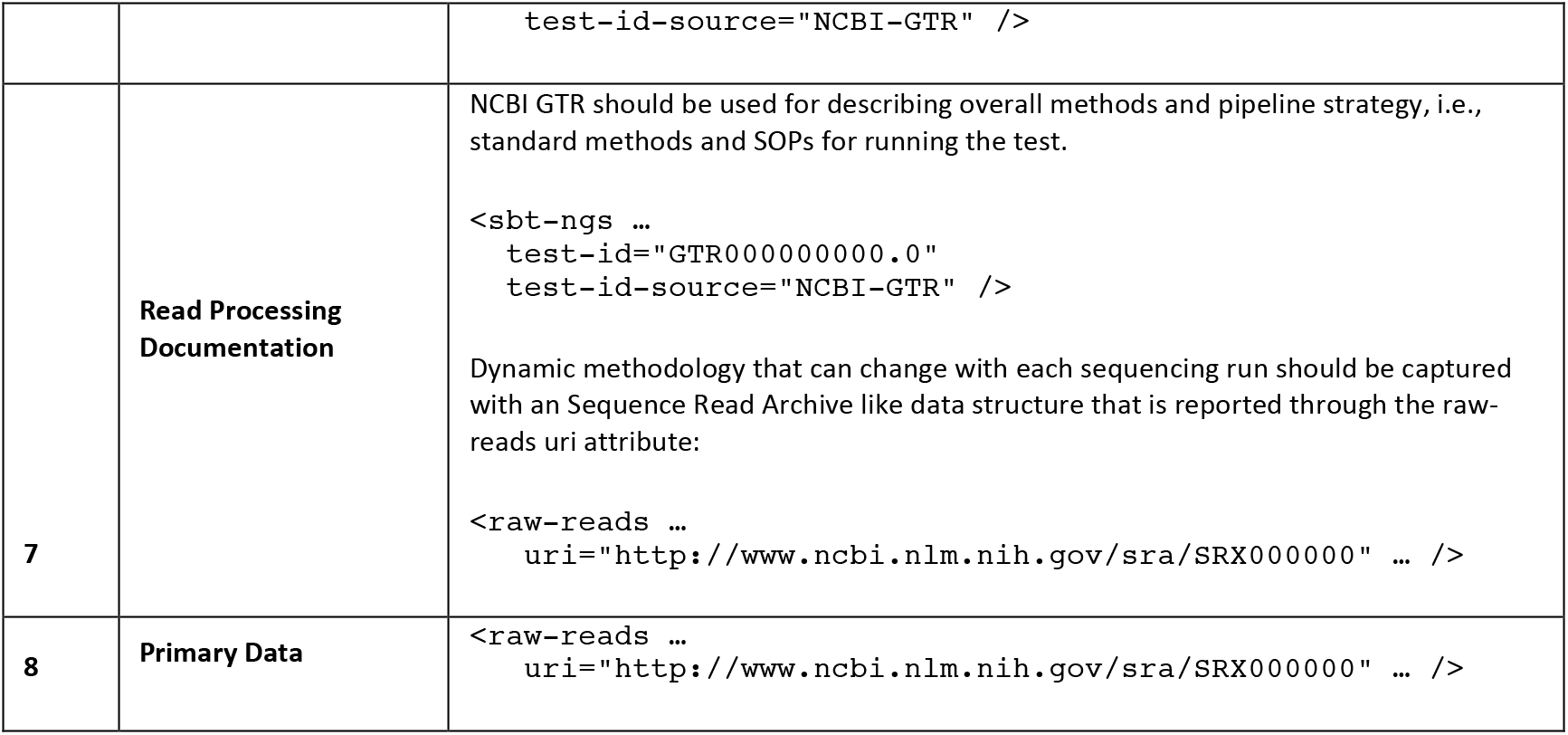
Mapping of MIRING elements to HML solutions

## 4.2. Validation

HML 1.0 has been finalized and the XML schema can be found on https://bioinformatics.bethematchclinical.org/HLA-Resources/HML/. Reporting of NGS based genotyping of HLA and KIR conforming to MIRING principles is possible using a combination of HML and semantic and syntactic validation tools, currently under development. A syntactically correct HML genotyping report may not pass organization specific business rules for acceptance or compliance to reporting guidelines such as MIRING. On the other hand, business-specific HML messages should conform to, and validate with, public XML schema definitions (XSDs). Transparent validation encourages a standards-based approach to mutual exchange of data for reproducible research and clinical application. Because of this, different validation tools will need to be developed depending on use cases and degree of semantic validation. For example, when a URI is presented to refer to external data and metadata, the validator will need to assess whether the link exists, and whether the information it points to makes sense and is interpretable. If HML is extended through the use of `property` elements and `anyAttribute` attributes, validation is further complicated since they are largely ignored with syntactic validation tools. Robust validation tools for these special use cases must then be created. A tool to validate whether a given instance of a HML message is MIRING compliant is being developed.

## 4.3. Future work

HML 1.0 has been developed to serve the needs of the immunogenomics community. However we recognize the need for interoperability with the larger healthcare community and the potential to interface with EMR systems. In light of this, the possibility to encapsulate HML in HL7 messages or clinical structured documents is being explored.

HML is a standard derived from the community members listed above. However, the NMDP has a commitment to using standards developed and maintained by other parties whenever possible. HL7 is an international standards development organization committed to the interchange of healthcare data. We are investigating the utility of OIDs (referred to above) as unique identifiers, as well as HL7 interchange formats such as Clinical Document Architecture (CDA) and Fast Healthcare Interoperability Resources (FHIR) to encapsulate HML. There is an active community to develop these standards that serves a larger audience than the parties listed in this document. We are actively engaged with the HL7 Clinical Genomics Working Group to ensure the content contained in the HML 1.0 specification can be contained in an HL7 message. This will allow the parties listed here to leverage the standards developed and implemented by the HL7 community. HL7 is also a party to the Biomedical Research Integrated Domain Group (BRIDG) as a consortium member including the FDA, CDISC, and NCI. The BRIDG members are developing artifacts to assist in the interchange of study-driven protocols and their regulatory artifacts, but also to assist in the interchange of molecular testing as part of the LSDAM (Life Sciences Domain Analysis Model). Since the HML 1.0 model is, at its core, an encapsulation of the results of a molecular test, we wish to leverage the work of those parties as well.

## 5. Acknowledgements

This work was supported by Office of Naval Research (ONR) grant N00014-12-1-0142 (MM and RPM), National Institutes of Health (NIH) grants U01AI067068 (SJM and JAH), awarded by the National Institute of Allergy and Infectious Disease (NIAID), and R01GM109030 (SJM, JAH, MM, and RPM), awarded by the National Institute of General Medical Sciences (NIGMS). The content presented is solely the responsibility of the authors and does not necessarily represent the official views of the NIH, NIAID, NIGMS, ONR, Department of Office of Naval Research, the Department of the Navy, the Department of Defense, or the US Government.

**Conflicts of Interest:** There are no conflicts of interest.

## References

[1] Erlich H. HLA DNA typing: past, present, and future. Tissue Antigens 2012;80:1–11.

[2] Robinson J, Halliwell JA, McWilliam H, Lopez R, Parham P, Marsh SGE. The IMGT/HLA Database. Nucleic Acids Res 2013;41:D1222–7.

[3] Voorter CEM, Mulkers E, Liebelt P, Sleyster E, van den Berg-Loonen EM. Reanalysis of sequence-based HLA-A, -B and -Cw typings: how ambiguous is today’s SBT typing tomorrow. Tissue Antigens 2007;70:383–9.

[4] Maiers M, Hurley CK, Perlee L, Fernandez-Vina M, Baisch J, Cook D, et al.. Maintaining updated DNA-based HLA assignments in the National Marrow Donor Program Bone Marrow Registry. Rev Immunogenet 2000;2:449–60.

[5] Hollenbach JA, Mack SJ, Gourraud P-A, Single RM, Maiers M, Middleton D, et al.. A community standard for immunogenomic data reporting and analysis: proposal for a STrengthening the REporting of Immunogenomic Studies statement. Tissue Antigens 2011;78:333–44.

[6] Helmberg W, Hegland J, Hurley CK, Maiers M, Marsh SG, Müller C, et al.. Going back to the roots: effective utilisation of HLA typing information for bone marrow registries requires full knowledge of the DNA sequences of the oligonucleotide reagents used in the testing. Tissue Antigens 2000;56:99–102.

[7] Mack SJ, Single RM, Erlich HA, Thomson G.. Proposal For HLA Data Validation. 2008. https://immport.niaid.nih.gov/docs/standards/Proposal For HLA Data Validation Version 2.doc, last accessed 29 January 2015

[8] Immunogenomic Next Generation Sequencing Data Consortium. http://ngs.immunogenomics.org last accessed 29 January 2015

[9] Mack SJ, Milius RP, Gifford BD, Sauter J, Hofmann J, Osoegawa K, et al.. Minimum Information for Reporting Next Generation Sequence Genotyping (MIRING): Guidelines for Reporting HLA and KIR Genotyping via Next Generation Sequencing. Hum Immunol 2015; **This issue**.

[10] Taylor CF, Field D, Sansone S-A, Aerts J, Apweiler R, Ashburner M, et al.. Promoting coherent minimum reporting guidelines for biological and biomedical investigations: the MIBBI project. Nat Biotechnol 2008;26:889–96.

[11] Maiers M.. A community standard XML message format for sequencing-based typing data. Tissue Antigens 2007;69:69–71.

[12] Milius RP, Mack SJ, Hollenbach JA, Pollack J, Heuer ML, Gragert L, et al.. Genotype List String: a grammar for describing HLA and KIR genotyping results in a text string. Tissue Antigens 2013;82:106–12.

[13] Data Standards Hackathon for NGS based genotyping. http://dash.immunogenomics.org, last accessed 29 January 2015

[14] HL7 Implementation Guidance for Unique Object Identifiers (OIDs), Release 1 2009:1-34. http://www.hl7.org/documentcenter/private/standards_temp_2E1D25F2-1C23-BA17-0C74CBDB29844F8B/v3/V3_OIDS_R1_INFORM_2011NOV.pdf, last accessed 29 January 2015

[15] Steindel SJ. OIDs: how can I express you? Let me count the ways. J Am Med Inform Assoc 2010;17:144–7.

[16] Rubinstein WS, Maglott DR, Lee JM, Kattman BL, Malheiro AJ, Ovetsky M, et al.. The NIH genetic testing registry: a new, centralized database of genetic tests to enable access to comprehensive information and improve transparency. Nucleic Acids Res 2013;41:D925–35.

[17] Gray KA, Yates B, Seal RL, Wright MW, Bruford EA. genenames.org: the HGNC resources in 2015. Nucleic Acids Res 2015;43:D1079–85.

[18] Danecek P, Auton A, Abecasis G, Albers CA, Banks E, DePristo MA, et al.. The variant call format and VCFtools. Bioinformatics 2011;27:2156–8.

[19] Terry SF. The Global Alliance for Genomics & Health. Genet Test Mol Biomarkers 2014;18:375–6.

[20] Global Alliance for Genomics & Health - Schemas for the Data Working Group. https://github.com/ga4gh/schemas, last accessed 29 January 2015

[21] James Kent W, Sugnet CW, Furey TS, Roskin KM, Pringle TH, Zahler AM, et al.. The human genome browser at UCSC. Genome Res 2002;12:996–1006.

[22] Cornish-Bowden A. Nomenclature for incompletely specified bases in nucleic acid sequences: recommendations 1984. Nucleic Acids Res 1985;13:3021–30.

[23] Eilbeck K, Lewis SE, Mungall CJ, Yandell M, Stein L, Durbin R, et al.. The Sequence Ontology: a tool for the unification of genome annotations. Genome Biol 2005;6:R44.

[24] Flanagan SE, Patch A-M, Ellard S. Using SIFT and PolyPhen to predict loss-of-function and gain-of-function mutations. Genet Test Mol Biomarkers 2010;14:533–7.

[25] Adzhubei I, Jordan DM, Sunyaev SR. Predicting functional effect of human missense mutations using PolyPhen-2. Curr Protoc Hum Genet 2013.

[26] Den Dunnen JT, Antonarakis SE. Mutation nomenclature extensions and suggestions to describe complex mutations: A discussion. Hum Mutat 2000;15:7–12.

[27] Human Genome Society recommendations for the description of sequence variants. http://www.hgvs.org/mutnomen/recs.html, last accessed 29 January 2015

